# Peripheral Skeletal Muscle Alterations in Adults Born Preterm: An Observational Comparative Study

**DOI:** 10.1101/2024.07.08.602584

**Authors:** Alyson Deprez, Ramy El-Jalbout, Anik Cloutier, Dany H. Gagnon, Andréa Gagnon Hamelin, Marie-Eve Mathieu, Thiffya A Kugathasan, Nicolas A. Dumont, Anne Monique Nuyt, Thuy Mai Luu

## Abstract

Prematurity is associated with reduced exercise capacity, which relies on the integrity of the cardiovascular, pulmonary, and skeletal muscle systems. Our animal model mimicking prematurity-associated conditions showed altered muscle composition and atrophy in adulthood. This study aimed to compare muscle composition and strength in adults born preterm versus full-term controls. This observational cohort study recruited 55 adults born preterm, ≤29 weeks’ of gestation and 53 full-term controls who underwent musculoskeletal ultrasound imaging to assess morphology of the rectus femoris at rest and during a maximal voluntary contraction. Maximal voluntary contraction of the hands and legs were measured by manual dynamometry. In adults born preterm, there was a reduction in muscle strength (handgrip: -4.8 kg, 95% CI -9.1, -0.6; knee extensor: -44.6 N/m, 95% CI -63.4, -25.8) and muscle area (-130 mm2, 95% CI -207, -53), which was more pronounced with a history of bronchopulmonary dysplasia. Muscle stiffness was increased in the preterm group (0.4 m/s, 95% CI 0.04, 0.7). Prematurity is associated with alterations in skeletal muscle composition, area, and function in adulthood. These findings highlight the necessity to implement preventive and/or curative approaches to improve muscle development and function following preterm birth to enhance overall health in this population.

**What’s known on This Subject:** Preterm birth is associated with reduced exercise capacity. However, the impact of preterm birth on skeletal muscle, a critical player of exercise capacity, in adulthood remains unclear.

**What This Study Adds:** Our findings provide novel insights into the potential long-term effects of preterm birth and the contributions of bronchopulmonary dysplasia on peripheral muscle-related health outcomes, such as muscle composition (reduced muscle area and increased muscle stiffness) and function (reduced muscle strength).

## Introduction

Improved perinatal care has allowed the survival of infants born extremely preterm (<28 weeks of gestational age (GA). As individuals born preterm reach adulthood, studies have shown their increased vulnerability for chronic health diseases (1). Indeed, preterm birth and its associated conditions can alter the normal sequence of organ development with lasting effects on future cardiovascular, pulmonary, or renal health (1). Moreover, adults born preterm who had complications such as bronchopulmonary dysplasia (BPD), the chronic lung disease of prematurity, are more susceptible to display organ system dysfunction in adulthood (2).

Several studies in children and adults born preterm reported reduced exercise capacity, mostly explained by cardiopulmonary impairment (3). Exercise capacity is also determined by the integrity of the skeletal muscle system, which is important to maintain posture, voluntary movements, and to support involuntary actions, such as breathing and protective reflexes (4). Skeletal muscles are further involved in organs and tissues regulation through the release of signaling molecules, the myokines (5, 6). Alteration of skeletal muscle tissue can disrupt homeostasis of several systems and contribute to heighten the risk for chronic diseases (7). The third trimester of pregnancy is an important period for skeletal muscle ontogenesis, characterized by maturation of muscle fiber and excitation-contraction coupling, and fiber type determination (8).

We previously conducted a systematic review and meta-analysis to summarize the impact of preterm birth on skeletal muscles and found a reduction of muscle thickness and power in individuals born preterm. However, studies remain scarce in adults born preterm with limited data on other characteristics of skeletal muscles, such as muscle area composition, and strength, highlighting the knowledge gap in understanding the effect of preterm birth on skeletal muscle development (9).

In a preclinical model mimicking deleterious conditions associated with preterm birth, preterm birth-related conditions cause an oxidative stress and inflammatory response associated with activation of protein degradation pathways, muscle atrophy, lower mitochondrial oxidative capacity, and muscle fatigability at juvenile and adult stages (10, 11).

We therefore postulated that preterm birth is associated with altered composition and function of the skeletal muscle in adults born very preterm which could be worse with a history of bronchopulmonary dysplasia. To tackle this hypothesis, we assessed skeletal muscle composition by musculoskeletal ultrasound of the *rectus femoris* and muscle strength using dynamometry in adults born preterm and compared their measures to full-term controls.

## Materials and methods

### Study population

The Health of Adults born Preterm Investigation (HAPI) cohort recruited individuals born at ≤29 weeks’ GA and full-term (≥37 weeks’ GA) controls. Participants were identified from a list of patients born at one of the three neonatal intensive care units in Montreal, Canada, between 1987-1997 (12). Full-term controls were born with birth weight ≥2500 grams and group matched for sex and age (±2 years). When available, we recruited them among friends and siblings of individuals born preterm, to take into consideration the shared environment. Exclusion criteria for all participants were the presence of severe neurosensory deficit preventing test completion or being pregnant (Supplemental figure 1). Of the 247 individuals reached, 109 were included. One participant was excluded due to gestational age (32 weeks). All participants gave written consent to participate in the study. They were assessed at Centre Hospitalier Universitaire Sainte-Justine from June 2021 to October 2023.

### Muscle composition

#### Image acquisition by ultrasound

B-mode ultrasound images of the rectus femoris (quadricep muscle) were captured with a Canon Aplio i800 device (Canon Medical Systems, Otawara-Shi, Japan) using a 14L5 linear probe set at 14 MHz to assess muscle area, thickness, echogenicity, and elasticity/stiffness. Images were acquired by two operators trained for high intra- and inter-observer reliability (ICC 0.94-0.99). Throughout the study, the gain was set at 75 dB (dynamic range of 75 gradations) and remained constant whereas the depth ranged between 5 and 7 cm to allow full visualization of the muscle and the focal zone was positioned at the middle of this muscle. All participants were in a sitting position with the knee in a 90-degree flexion. Images were taken from the non dominant leg (defined as the opposite leg use to kick a ball or climb stairs) halfway along the line from the anterior superior iliac spine to the superior border of the patella. Three B-mode images were taken for each plane: transverse and longitudinal at rest and under maximal voluntary contraction. The elasticity/stiffness of the rectus femoris was also evaluated in triplicates using single shot shear wave elastography (SWE) at rest and under maximal voluntary contraction in a longitudinal plane.

#### Image analysis

Muscle area (cross sectional area, mm^2^) was assessed using images captured in the transverse plane. Sub-cutaneous thickness (cm) (distance between the skin and the superior fascia of the muscle), echogenicity (average pixel value in a region of interest, expressed in arbitrary unit - AU), and shear wave elastography (m/s) were assessed using images captured in the longitudinal plane. Muscle area and echogenicity analyses were performed by a single operator using a custom program (USLBP_GUI; version R2018b) developed with MATLAB’s Image Processing Toolbox (The MathWorks Inc., Natick, MA, USA), as used in previous studies (Supplemental figure 2) (13) Muscle area and thickness were normalized to the body mass index (BMI)(14).

### Handgrip and quadriceps strength

Grip strength was assessed with participants seated, both feet flat on the ground. Arm was unsupported with elbow flexion at 90 degrees. Participants were encouraged to squeeze the handle of the dynamometer (Lafayette Hand Dynamometer, model 78010) to generate a maximal voluntary effort held during at least three seconds, for three measures for both hands. Maximum values obtained on the dominant and non-dominant hand were averaged to calculate the mean maximal grip strength. The absolute maximal grip strength out of all six trials was also reported (15, 16). Values are reported as kilograms (kg) and normalized to height (m) (17). For knee extensor strength, an instrumented dynamometer (EasyForce, Meloq AB, Sweden) was fixed from the table to the ankle of the dominant and non-dominant legs with the knee and hip flexed at 90 degrees with participants seated. After three submaximal trials to warm up, participants performed three maximal isometric voluntary contractions on the dominant and non-dominant legs with a one-minute rest period between each measurement. Maximum values from each leg were averaged to calculate legs extensor strength. Values are reported in Newton meters (Nm) and normalized to body weight (kg) (18). For normalization of the strength to muscle surface area (cm^2^) (evaluated by ultrasound), value from the non-dominant leg was used.

### Additional assessments

Height, weight, thigh circumference, body mass index, daily count step, physical activity, estimated VO_2_ max (based on Huet questionnaire)(19) and neonatal characteristics are described in the online supplementary material.

### Statistical analysis

Descriptive statistics were calculated as mean with standard deviation (SD) for continuous variables and counts with proportions for categorical variables. All between-group comparisons were performed using linear regression analyses and adjusted for sex. Given the ultrasound absorbance properties of adipose tissue, we further adjusted for sub-cutaneous thickness when comparing muscle echogenicity. All analyses were performed using SPSS software version 27 and the graphical representations of correlations were done with R software.

## RESULTS

### Study population

We included 108 participants: 55 very preterm (24 men, 31 women; mean GA: 26.9±1.3 weeks) and 53 full-term controls (23 males, 30 women; mean GA 39.4±1.3 weeks) (Table 1). Among adults born preterm, participants were of lower GA (non-participants: 27.2±1.3 weeks, P=0.383) and birthweight (non-participants: 1033±274 grams, P=0.045), but there was no sex difference (non-participants: 55% females, P=0.940). Mean age at assessment was 29.9 years. In the preterm group, 16 (29%) had a history of moderate to severe BPD. In addition, 5 had some level of neuromotor deficits (i.e., cerebral palsy), but were still able to complete the study protocol. Individuals born preterm had smaller anthropometric measures compared to full-term controls. Estimated VO_2_max [ml/(kg/min)] was lower in the preterm group. There were no observed between-group differences in the number of steps, sedentary time, moderate-to-vigorous and vigorous physical activity time per day.

**Table 1:**
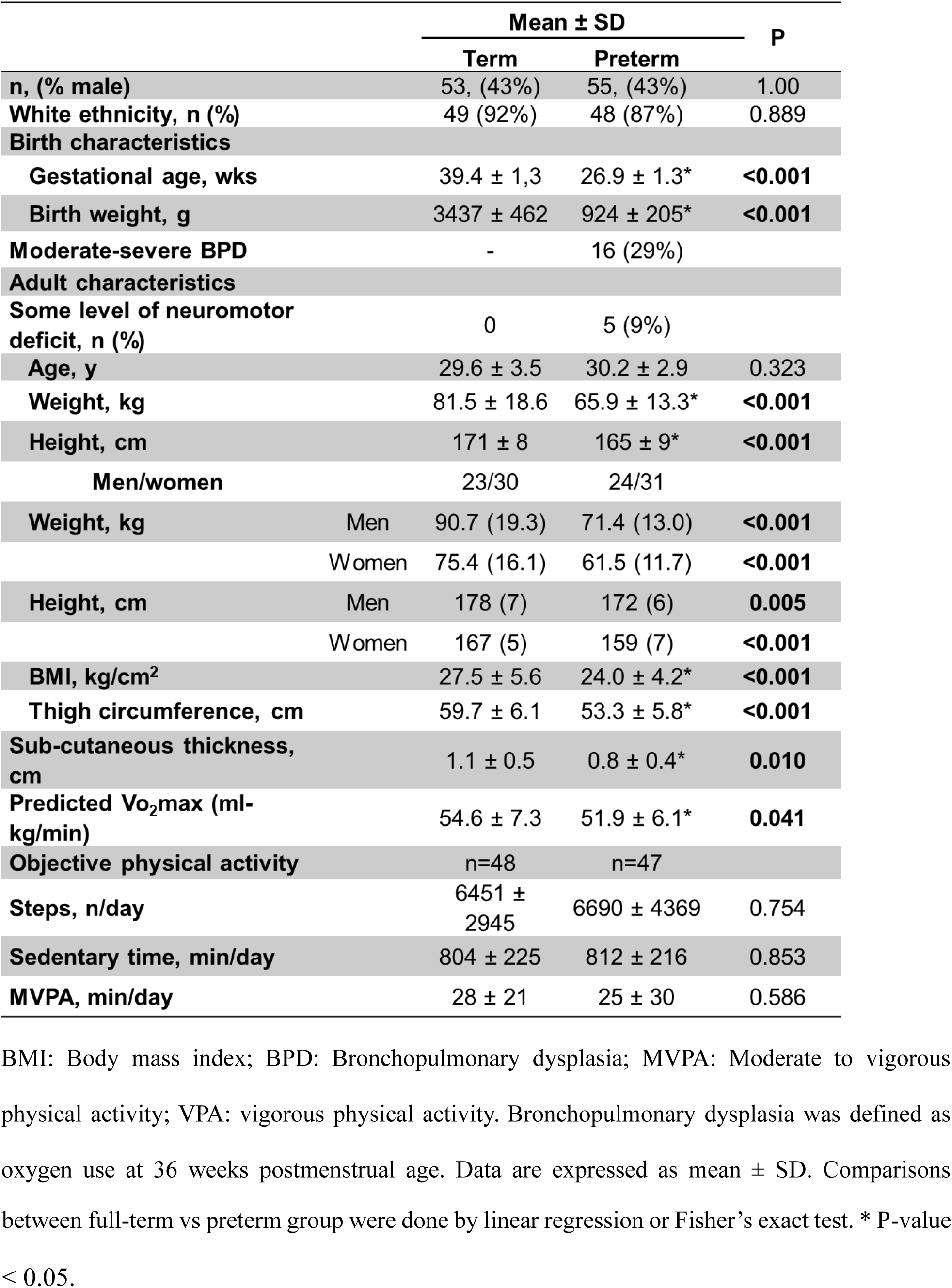
Population characteristics.

### Muscle structure

Individuals born preterm had a reduction of 13.5 % of the muscle area at rest (mean difference, MD: of -130 mm^2^; 95% CI: -206 to -54, P =0.001) and 12.5 % during maximal contraction (MD: - 119 mm^2^; 95% CI -202 to -35, P=0.006) (Table 2 and Supplemental figure 3) even after adjusting for sex. However, when normalizing to BMI, differences of muscle area and thickness between individuals born preterm and full-term were no longer observed. Muscle stiffness (shear wave elastography, SWE) was higher in adults born preterm during maximal contraction adjusting for sex (MD: +0.4 m/s; 95% CI 0.0 to 0.7, P=0.026). A similar trend was observed at rest, although it did not reach statistical significance. No between-group differences in muscle echogenicity were found at rest or at maximal contraction (Table 2).

**Table 2:** Muscle composition.

|  | Mean (SD) |  | Mean difference<br>(95% CI) | P | Adjusted mean<br>difference (95% CI) | P |
| --- | --- | --- | --- | --- | --- | --- |
|  | Term<br>n=53 | Preterm<br>n=55 |  |  |  |  |
| *At rest* |  |  |  |  |  |  |
| Muscle area, mm<sup>2</sup> | 966 (219) | 835 (182) | -130 (-207, -53) | **0.001** | -130 (-206, -54)* | **<0.001** |
| Muscle area, mm<sup>2</sup>/BMI | 36.1 (9.3) | 35.3 (8.2) | -0.7 (-4.1, 2.5) | 0.655 | 0.7 (-4.1, 2.5) | 0.650 |
| Muscle thickness, mm | 19.6 (3.8) | 18.1 (3.3) | -1.5 (-2.8, -0.1) | **0.031** | -1.5 (-2.8, 0.1)* | **0.028** |
| Muscle thickness, mm/BMI | 0.73 (0.17) | 0.76 (0.14) | 0.03 (-0.02, 0.09) | 0.294 | 0.03 (-0.02, 0.09) | 0.293 |
| Shear wave elastography, m/s | 2.3 (0.2) | 2.4 (0.3) | 0.1(-0.0, 0.2) | 0.067 | 0.1 (-0.0, 0.2)+ | 0.066 |
| Echogenicity, AU | 53.5 (8.7) | 56.1 (8.2) | 2.6 (-0.5, 5.9) | 0.106 | 0.6 (-2.5, 3.9)† | 0.672 |
| *Maximal contraction* |  |  |  |  |  |  |
| Muscle area, mm<sup>2</sup> | 948 (240) | 829 (207) | -118 (-204, -33) | **0.007** | -119 (-202, -35)* | **0.006** |
| Muscle area, mm<sup>2</sup>/BMI | 35.3 (9.7) | 35.2 (9.6) | -0.1 (-3.8, 3.5) | 0.922 | 0.1 (-3.8, 3.4) | 0.918 |
| Muscle thickness, mm | 23.2 (4.0) | 22.3 (3.7) | -0.9 (-2.4, 0.5) | 0.223 | -0.9 (-2.4, 0.5)* | 0.218 |
| Muscle thickness, mm/BMI | 0.87 (0.19) | 0.95 (0.20) | 0.08 (0.00, 0.15) | **0.040** | 0.08 (0.03, 0.15) | **0.040** |
| Shear wave elastography, m/s | 2.7 (0.8) | 3.1 (1.0) | 0.4 (0.04, 0.7) | **0.027** | 0.4 (0.0, 0.7) + | **0.026** |
| Echogenicity, AU | 50.3 (10.9) | 53.5 (9.9) | 3.2 (-0.7, 7.2) | 0.111 | 0.4 (-3.4, 4.2) † | 0.830 |

AU= arbitrary unit. Data are expressed as mean ± SD. Muscle area (mm<sup>2</sup>) and thickness (mm) are normalized by the body mass index (BMI). Comparisons between full-term vs preterm group were done by linear regression adjusted for sex \*, sex and subcutaneous tissue thickness †; P values are reported in the table.

### Muscle function

Individuals born preterm, compared to full-term controls, had reduced maximal knee extensor strength (MD: -44.7 Nm; 95% CI -61.4, -28.0, P<0.001), even after normalizing for body weight and accounting for sex (Table 3). Combined and absolute maximal handgrip were lower for the preterm group in comparison to their full-term counterparts (Table 4) (MD: -4.8 kg; 95% CI -9.1, -0.6, P=0.025 and -4.3 kg; 95% CI -8.8, 0.1, P= 0.058, respectively). When stratified by sex, the preterm-term differences were larger among men than among women.

**Table 3:** Leg extensor strength.

|  | Term<br>n=53 | Preterm<br>n=55 | Mean difference<br>(95% CI) | *P* | Adjusted mean<br>difference (95% CI) | *P* |
| --- | --- | --- | --- | --- | --- | --- |
| **Leg extensor strength, Nm** | 133.5 (56.2) | 88.9 (41.4) | -44.6 (-63.4, -25.8) | **<0.001** | -44.7 (-61.4, -28.0) | **<0.001** |
| **Leg extensor strength, Nm/kg** | 4.2 (1.5) | 3.7 (1.3) | -0.5 (-1.1, -0.0) | **0.046** | -0.6 (-1.1, -0.0) | **0.037** |
| **Leg extensor strength, Nm/cm<sup>2</sup>** | 0.13 (0.06) | 0.10 (0.04) | -0.03 (-0.05, -0.01) | **0.002** | -0.03 (-0.04, -0.00) | **<0.001** |

Data are expressed as mean ± SD. Comparisons between full-term vs preterm group were done by linear regression and adjusted for sex; *P* values are reported in the table. Leg extensor strength Nm and Nm/kg are the average of the maximum values obtained from the non-dominant and dominant legs. Leg extensor strength Nm/cm<sup>2</sup> is the maximum value from the non-dominant leg normalized to the muscle surface area from ultrasound analysis.

**Table 4:** Handgrip strength.

|  | Sex | Mean (SD) |  | Mean difference<br>(95% CI) | P |
| --- | --- | --- | --- | --- | --- |
|  |  | Term<br>n=49 | Preterm<br>n=54 |  |  |
| **Combined maximal handgrip, kg** |  | 39.9 (11.2) | 35.0 (10.4) | -4.8 (-9.1, -0.6) | **0.025** |
| **Absolute maximal handgrip, kg** |  | 42.1 (11.5) | 37.7 (11.3) | -4.3 (-8.8, 0.1) | 0.058 |
| **Men/women** |  | 20/29 | 23/31 |  |  |
| **Combined maximal handgrip, kg** | Men | 51.3 (7.9) | 43.6 (8.4) | -7.6 (-12.7, -2.5) | **0.004** |
|  | Women | 32.1 (4.4) | 28.7 (6.4) | -3.4 (-6.3, -0.5) | **0.021** |
| **Absolute maximal handgrip, kg** | Men | 53.9 (7.9) | 46.5 (8.6) | -7.3 (-12.4, -2.1) | **0.006** |
|  | Women | 33.9 (4.3) | 31.2 (8.4) | -2.7 (-6.2, -0.7) | 0.122 |
| **Combined maximal handgrip, kg/m** | Men | 28.8 (4.1) | 25.1 (4.0) | -3.7 (-6.2, -1.1) | **0.005** |
|  | Women | 19.2 (2.5) | 17.9 (3.8) | -1.3 (-3.0, 0.3) | 0.127 |
| **Absolute maximal handgrip, kg/m** | Men | 30.2 (4.0) | 26.8 (4.3) | -3.4 (-6.0, -0.8) | **0.010** |
|  | Women | 20.3 (2.4) | 19.5 (5.0) | -0.8 (-2.9, 1.2) | 0.420 |

Data are expressed as mean ± SD. Comparisons between full-term vs preterm group were done by linear regression; P values are reported in the table.

### Bronchopulmonary dysplasia and skeletal muscle

To explore factors that could contribute to skeletal muscle alterations, we compared skeletal muscle composition and function between adults born preterm with moderate to severe BPD and those without BPD. Muscle area at rest was reduced in those with BPD (MD: -119 mm^2^, 95% CI: -235, -3, P=0.044) even after adjusting for birth weight. We did not identify any statistically significant differences for the other parameters (Table 5).

**Table 5:** Comparison of muscle composition and strength between individuals born preterm with and without bronchopulmonary dysplasia (BPD).

|  | Mean (SD) |  | Mean difference<br>(95% CI) | P | Adjusted mean<br>difference (95%<br>CI) | P |
| --- | --- | --- | --- | --- | --- | --- |
|  | Preterm – No<br>BPD | Preterm –<br>BPD |  |  |  |  |
|  | n=39 | n=16 |  |  |  |  |
| At rest |  |  |  |  |  |  |
| Muscle area, mm<sup>2</sup> | 867 (181) | 744 (153) | -122 (-226, -19) | 0.021 | -119 (-235,-3) | 0.044 |
| Shear wave elastography,<br>m/s | 2.4 (0.3) | 2.5 (0.5) | 0.1 (-0.1, 0.3) | 0.267 | 0.1 (-0.0, 0.4) | 0.204 |
| Maximal contraction |  |  |  |  |  |  |
| Muscle area, mm<sup>2</sup> | 854 (217) | 768 (172) | -85 (-208, 36) | 0.165 | -83 (-220, 53) | 0.226 |
| Shear wave elastography,<br>m/s | 3.0 (1.0) | 3.4 (0.9) | 0.4 (-0.2, 1.1) | 0.178 | 0.5 (-0.1, 1.2) | 0.115 |
| Muscle function | n=38 | n=16 |  |  |  |  |
| Handgrip, kg | 34.7 (10.1) | 35.8 (11.3) | 1.0 (-5.2, 7.3) | 0.746 | 4.8 (-1.8, 11.4) | 0.152 |
| Leg extensor strength, Nm | 92.2 (45.0) | 74.3 (30.5) | -17.8 (-42.5, 6.8) | 0.152 | -9.8 (-36.4, 16.7) | 0.461 |

Bronchopulmonary dysplasia (BPD) was defined as oxygen use at 36 weeks postmenstrual age.
Data are expressed as mean ± SD. Comparisons between No BPD vs BPD group were done by linear regression and adjusted for birthweight; P values are reported in the table.

### Relationship between skeletal muscle strength, muscle composition and aerobic capacity

Individuals with greater skeletal muscle mass at the quadriceps typically displayed greater knee extensor strength. For each cm^2^ increase in cross-sectional area of the quadriceps, knee extensor strength increased by 9.4 Nm (95% CI 3.7, 15.1) in the preterm group and by 10.4 Nm (95% CI 3.8, 16.9) in the full-term group (Figure 1A). Conversely, increased skeletal muscle stiffness was associated with reduced strength. Each unit increase in SWE resulted in a decrease in knee extensor strength of -18.2 Nm (-27.8, -8.7) and -25.6 Nm (-39.9, -11.4) for the preterm and full-term born groups, respectively (Figure 1B). Finally, predicted VO_2_ max was associated with muscle strength in both groups (Supplemental Table 1). However, we did not find any association between level of physical activity and muscle area and strength (Supplemental Table 1).

**Figure 1:**
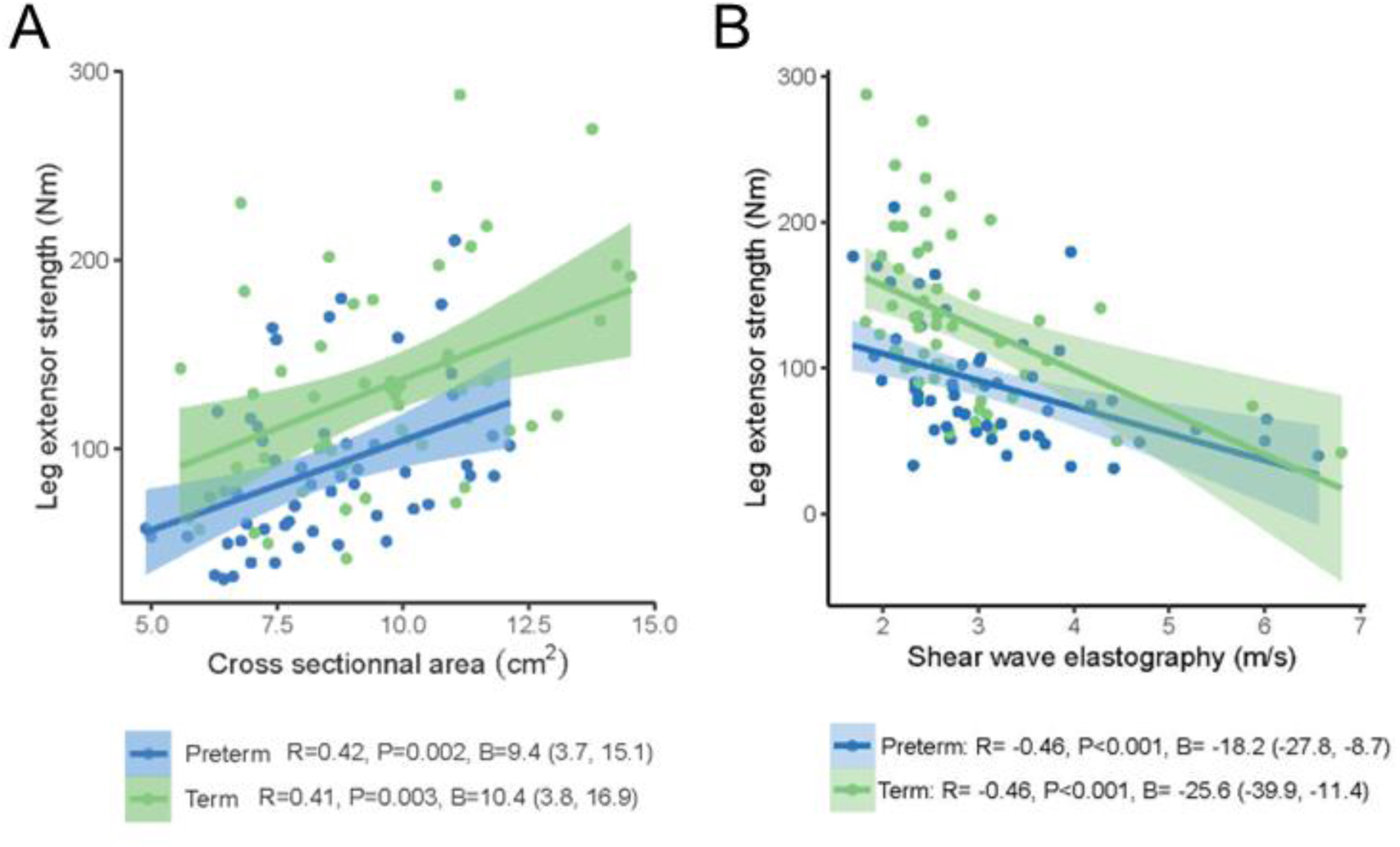
Association between cross sectional area, shear wave speed and knee extension strength. (A) Increased cross sectional area correlates with higher strength. (B) Increased shear wave elastography is associated with reduced strength. A linear regression line is displayed for the association between the two variables with its computed 95% CI. Correlation coefficients (R) and P-values were calculated using the Pearson’s method.

## DISCUSSION

This study shows that adults born very preterm have altered skeletal muscle health. Compared to full-term controls, they have reduced muscle mass, even more so with a history of BPD, reduced strength, and increased muscle stiffness. However, there is no significant difference in muscle echogenicity and level of physical activity among the participants. Increased muscle mass and lower stiffness are associated with greater strength, the latter also correlating with predicted VO_2_max. To our knowledge, very few studies have concomitantly investigated muscle composition and strength in adults born very preterm. None has evaluated muscle stiffness nor the effect of BPD on skeletal muscle health.

Skeletal muscle health is an important determinant of functioning and quality of life. In older individuals, sarcopenia (i.e., the loss of skeletal muscle mass and strength) is associated with a reduced capacity to perform activities of daily living (20). The loss of function can result in lower mobility and frailty, but also metabolic problems, which, in turn, can predispose to adverse health conditions and premature death (21). Preterm birth is associated with reduced exercise capacity, glucose intolerance, and increased risk of cardiovascular diseases (1, 3, 22, 23). The extent to which the skeletal muscle could contribute to these dysfunctions is unknown. The first step was therefore to examine whether skeletal muscle tissue was altered in adults born preterm, which could potentially point towards a target for intervention to improve health.

Preterm birth can disrupt several maturational processes important to healthy organ development, including the skeletal muscle tissue (24). Mechanisms are still under investigations, but could involve early life exposure to oxidative stress (25, 26) and systemic inflammation (27). Complications like BPD or sepsis, and treatments, including steroids or anti-inflammatory agents, could also influence skeletal muscle tissue development (28, 29). In a preclinical rodent model mimicking preterm birth-related conditions, markers of inflammation and oxidative stress were found within the skeletal muscle tissue of juvenile pups exposed to transient neonatal hyperoxia. These exposed tissues displayed reduced fiber size, increased proportion of fast fatigable fibers, and increased deposition of collagen over time and decreased muscle strength, which was more pronounced in males than females (10, 11). This regulatory axis between inflammation and muscle has also been observed in other conditions. For instance, low-grade systemic inflammation has been associated with muscle atrophy in chronic obstructive pulmonary disease (COPD) and aging (30, 31). Muscle biopsies in patients with COPD have further shown upregulation of pro-inflammatory cytokines (32–34) along with muscle weakness.

In infants born preterm, muscle thickness was reduced compared to full-term controls (35), and even more so with BPD (36). We found similar findings, suggesting that smaller muscle size may track from infancy to adulthood. In addition, we observed that decreased muscle area was associated with reduced muscle strength of the quadriceps. The smaller height and weight, which characterize the body phenotype of individuals born preterm, could explain why muscle area and thickness are reduced. Indeed, when we adjusted for body size, the observed differences were no longer statistically significant. However, leg strength remained lower in the preterm group, in agreement with other studies (37–39), suggesting that there may be other contributors beside anthropometric differences. Lower leg strength could indicate altered myofibers function due to inadequate excitation-contraction coupling and/or ATP supply secondary to mitochondrial deficiency. In humans, autopsy studies found reduced mitochondria energy metabolism in the muscles of newborns born very preterm versus full-term (40, 41). In a rodent model mimicking preterm birth-related conditions, it was showed a lower muscle mitochondrial biogenesis and oxidative activity, and compensatory higher glycolytic enzyme expression compared to control rats; these differences were observed in males but not in female rats (10) (37–39). We also found reduced strength in the upper extremities for the preterm group with a handgrip estimated to be equivalent to that of a 65-year-old male and a 60-year-old female (42). This corroborates results from Morrison and colleagues on 95 adults in their thirties with birth weight <1000 g, who displayed an absolute maximal grip strength comparable to individuals aged 65 years for males and 55-60 years for females (15), suggesting premature muscle wasting. Adult males born preterm appear more affected than females. Male infants born preterm are more vulnerable than females with higher mortality and morbidities (43–45). Some of these morbidities involve injury associated with oxidative stress and inflammation (46, 47). Female neonates versus males have greater antioxidant capacity (48, 49), which may confer better protection.

We assessed muscle stiffness/elasticity using elastography imaging. Measurement of muscle elasticity at rest provides relevant baseline physiological characteristics. However, measuring stiffness during muscle contraction is superior at discriminating a normal from a pathological state (50, 51). In our study, we observed at rest a slight increase of SWE, and significant differences with full-term controls during maximal contraction. Interestingly, increased stiffness was associated with reduced muscular strength. Considering that muscle stiffness increases with age (52, 53), longitudinal follow-up will allow to determine whether muscle aging occurs more prematurely and/or is accelerated in preterm population.

Physical activity is a critical determinant of skeletal muscle health. Cohort studies have shown that adults born very preterm self-report decrease leisure time physical activities compared to full-term peers (54). We objectively quantified physical activity and sedentary time using accelerometers and found no between-group differences, like others (55–57), suggesting that the observed skeletal muscle dysfunction is not solely the result of deconditioning. Moreover, these findings indicate that the reduction observed in the estimated aerobic capacity and the muscular strength are not a limit to physical activity in early adulthood. However, we could hypothesize that performing the same physical activity will require more efforts in preterm born individuals. The estimated aerobic capacity was surprisingly high, especially when considering both groups’ modest levels of physical activity. In previous studies, estimated VO_2_max using the Huet questionnaire was higher in our participants when compared to measured peak VO_2_ by cardiopulmonary exercise testing (58, 59). Nevertheless, estimated and measured VO_2_max are highly correlated (19), and the magnitude of the between-group differences using estimated values was comparable to what was previously reported with measured peak VO_2_ (12, 59).

Study limitations must be acknowledged. First, selection bias may have resulted in recruiting adults born preterm of higher socio-economic status and in better health. This would lead to an underestimate of the effect of preterm birth on skeletal muscle health. However, participants born preterm compared to non-participants were of smaller gestational age and lower birthweight, which could result in an overestimation of the difference. In addition, our cohort was predominantly white and representative of the general population in the province of Québec, Canada, born during that time period. Replication in other populations is required for generalizability. A portable electronic instrumented dynamometer was used to maximize the feasibility and generalizability; however, it comes inherently with limits if compared with a high-tech motorized instrumented dynamometer that would have provided added stability and reduced risk of movement compensation. This being said, a standardized protocol was followed in the present study as described above throughout the study using the same table and foam cushions for adjustment. Participants were familiarized to the testing procedure prior to performing the measures. Finally, the use of a questionnaire to estimate VO_2_max is not as objective as a direct measure. Future studies would gain in precision measuring VO_2_max.

## CONCLUSIONS

Our findings indicate that very preterm birth leads to alteration in skeletal muscle composition and function in adulthood. These alterations could contribute to the higher risk of developing cardiopulmonary disorders observed in preterm born individuals and hinder maintenance of an active lifestyle with aging. Interventions to improve skeletal muscle health may represent a novel avenue to prevent chronic health diseases following preterm birth. Physical activity can reverse aging in sarcopenic patient (60); an adapted exercise program targeting muscle mass and function from the neonatal intensive care unit to adulthood could bring benefits.

## Acknowledgments

We thank Hakim Mecheri for contributing to ultrasound images analyses, Dr Alexis Vivoli for the graphical generation on R software, and Mi-Suk Dufour for her helpful review of the statistical analysis. We thank all the participants for their contribution to this study. We thank the Meloq AB Company for kindly providing us the dynamometer used in this study.

## Author’s contributions

Alyson Deprez, Ramy El-Jalbout, Dany Gagnon, Nicolas A. Dumont, Anne Monique Nuyt and Thuy Mai Luu conceptualized and designed the study. Alyson Deprez, Ramy El-Jalbout, Andréa Gagnon-Hamelin, Anik Cloutier, Marie-Eve Mathieu and Thiffya A. Kugathasan designed the data collection instruments, collected data, carried out the initial analyses. Alyson Deprez drafted the initial manuscript and revised the manuscript. Thuy Mai Luu, Anne Monique Nuyt and Nicolas A. Dumont coordinated the project and supervised data collection. Ramy El-Jalbout, Dany H. Gagnon, Marie-Eve Mathieu, Nicolas A. Dumont, Anne Monique Nuyt and Thuy Mai Luu critically reviewed and revised the manuscript. All authors approved the final manuscript as submitted and agree to be accountable for all aspects of the work.

## Ethical guidelines

All participants have been approved by the appropriate ethics committee and have therefore been performed in accordance with the ethical standards laid down in the 1964 Declaration of Helsinki and its later amendments. All participants gave their informed consent prior to their inclusion in the study.

## Data availability statement

Data is available for sharing upon reasonable request to the corresponding authors.

