## Supplementary material for "Peripheral Skeletal Muscle Alterations in Adults Born Preterm: An Observational Comparative Study": online supplemental data

**Affiliations:** [1] Research Center, CHU Sainte-Justine; [2] Department of pharmacology and physiology, Faculty of Medicine, Université de Montréal, Canada; [3] School of Rehabilitation, Faculty of Medicine, Université de Montréal, Canada; [4] School of Kinesiology and Physical Activity Science, Faculty of Medicine, Université de Montréal, Canada; [5] Department of pediatrics, Faculty of Medicine, Université de Montréal, Canada

**Corresponding author:** Dr. Thuy Mai Luu, Sainte-Justine University Hospital and Research Center, 3175 Chemin de la Côte-Sainte-Catherine, Montréal (Québec) H3T 1C5 Canada. Tel.: +1 514 345 4931, ext 6642,

**ONLINE SUPPLEMENT**

### Materials and methods

#### Other health assessment

On the day of the study visit, height, weight, and thigh circumference were measured in triplicates. Body mass index, expressed in units of $kg/m^2$, was calculated by dividing the body mass (kg) by the square of the body height (m).

Participants completed the Huet questionnaire, a validated 10-item self-reported assessment of aerobic capacity, to estimate maximal oxygen consumption ($VO_2max$) (1). In addition, participants were asked to wear an ActiGraph flagship activity monitor (wGT3X-BT) for 7 days. Data were deemed valid if the device was worn for at least 10 hours per day on 4 weekdays and one weekend day. The epoch length was 60 seconds, and the non-wear time was defined by an interval of at least 60 consecutive minutes of zero activity intensity counts. Data were evaluated by Zero count program using Matlab software (2).

#### Neonatal characteristics

Neonatal data were collected from medical charts and/or immunization booklet, which contains information on gestational age, birth weight, APGAR score and age at discharge. Moderate to severe bronchopulmonary dysplasia (BPD) was defined as supplemental oxygen use after 36 weeks post-menstrual age (3).

**Results**

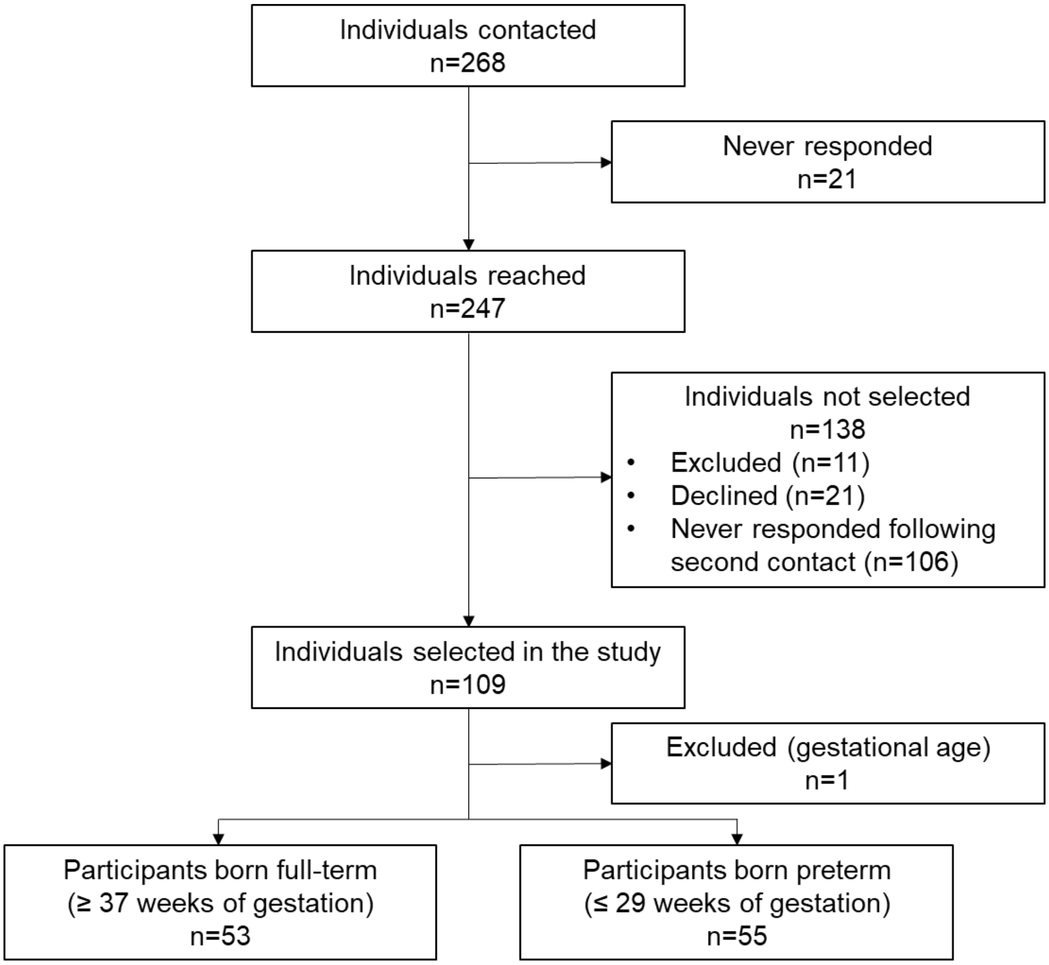


**Supplemental figure 1: Flow chart of the study**

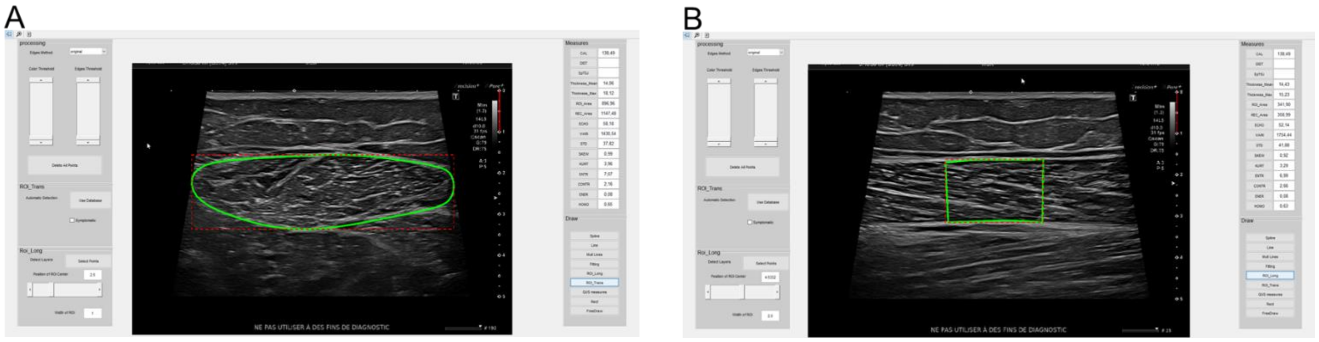


**Supplemental figure 2: Analysis of ultrasound images in the Matlab software.**

Representative ultrasound images of the analysis using Matlab software in transversal (A) and longitudinal (B) plane.

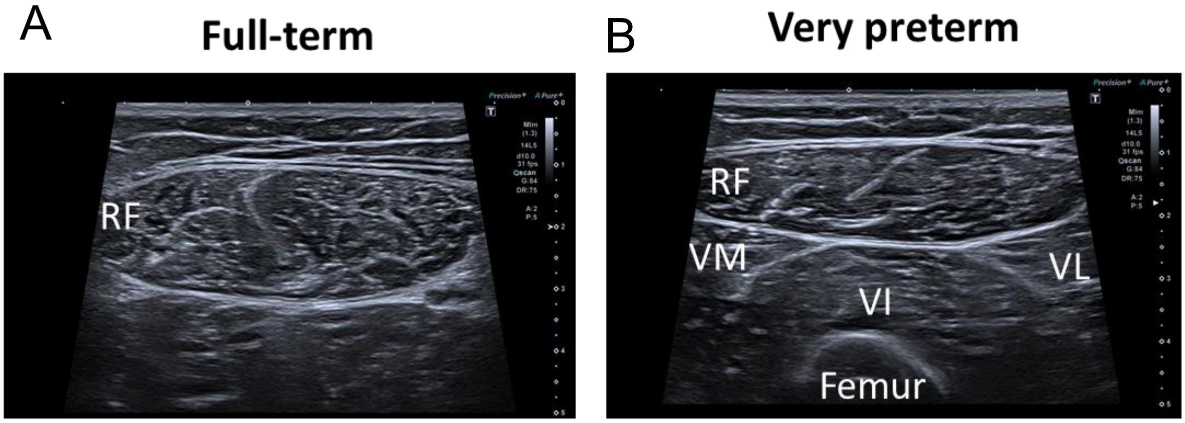


**Supplemental figure 3: Ultrasound images.** Representative ultrasound images in the transverse plane of the rectus femoris from individual born full-term (A) and very preterm (B). RF= Rectus Femoris; VI= Vastus Intermedius; VL= Vastus Lateralis; VM= Vastus Medialis.

**Supplemental Table 1: Association between muscle parameters and predicted $VO_2$ max and moderate to vigorous physical activity (MVPA).**

| | B Coefficients (95% Confidence Interval) | |
|---|---|---|
| | **Predicted $VO_2$max ml/(kg*min)** | **MVPA, min/day** |
| **Term** | *n=53* | *n=48* |
| Muscle area | 7.3 (-0.7, 15.3) | 1.9 (-0.9, 4.8) |
| Muscle strength | **2.1 (0.1, 4.0)** * | 0.2 (-0.4, 1.0) |
| **Preterm** | *n=55* | *n=48* |
| Muscle area | 5.6 (-2.1, 13.4) | 1.2 (-0.4, 2.8) |
| Muscle strength | **3.1 (1.7, 4.5)** * | 0.6 (-0.3, 0.7) |

Data are expressed as mean ± SD. Comparisons by linear regression, *P<0.05.
